# *Tetrahymena thermophila* Granule Lattice Protein 3 Improves Solubility of Sexual Stage Malaria Antigens Expressed in *Escherichia coli*

**DOI:** 10.1101/2021.06.29.450425

**Authors:** Cengiz Akkale, Donna Marie Cassidy-Hanley, Theodore G. Clark

**Affiliations:** Department of Mıcrobiology and Immunology, College of Veterinary Medicine, Cornell University, Ithaca, NY, USA; Department of Bioengineering, Faculty of Engineering, Adana Alparslan Türkeş Science and Technology University, Adana, Turkey

**Author notes:** **Correspondence**: Cengiz Akkale.

**Keywords:** malaria, vaccine, transmission blocking, *Tetrahymena thermophila*, granule lattice protein

## Abstract

The requirement for low cost manufacturing makes bacterial cells a logical platform for the production of recombinant subunit vaccines for malaria. However, protein solubility has been a major stumbling block with prokaryotic expression systems. Notable examples include the transmission blocking vaccine candidates, Pfs25 and Pfs48/45, which are almost entirely insoluble when expressed as recombinant proteins in *Escherichia coli*. Various solubility tags have been used with limited success in improving solubility, although recent studies with granule lattice protein 1 (Grl1p) from the ciliated protozoan, *Tetrahymena thermophila*, have shown promise. Here, we examine a related solubility tag, granule lattice protein 3 (Grl3p) from *T. thermophila*, and compare it to both Grl1p and the well-studied maltose binding protein (MBP) used to improve the solubility of multiple protein targets. We find that Grl3p performs comparably to Grl1p when linked to Pfs25 but significantly improves solubility when paired with Pfs48/45.

## 1. Introduction

Approximately half of the world’s population lives under the threat of malaria (Borchert et al., 2015). According to 2018 World Malaria Report data, ∼ 228 million clinical cases of the disease were reported worldwide, of which 405,000 resulted in death. *Plasmodium falciparum*, one of the five malarial species that infect humans, is responsible for the highest levels of morbidity and mortality (WHO, 2018).

Despite decades of effort and massive investment from both government agencies and the pharmaceutical industry, effective subunit vaccines for malaria have been largely unsuccessful with the possible exception of the recently tested R21/Matrix-M vaccine (Datoo et al., 2021). Given that natural immunity against disease often takes years to develop even with repeated exposure to infected mosquitos, vaccine failures are not entirely surprising. Following entry into the skin, sporozoite-stage parasites migrate rapidly to the liver where they infect hepatocytes and undergo schizogony to form tens of thousands of liver-stage merozites. Merozoites then emerge, enter the vasculature and infect red blood cells where they continue to expand, eventually causing clinical disease. Each stage of the parasite life cycle is accompanied by different sets of antigens, which themselves can undergo considerable variation. Combined with rapid intracellular division, antigenic variation provides an effective strategy of immune evasion and has limited the utility of vaccines targeting subunit antigens expressed in the human host (Greenwood et al., 2008).

By contrast, variation among parasite antigens expressed in the mosquito vector appears to be constrained and vaccine strategies that can interfere with malarial development in the mosquito have the potential to block transmission entirely and eradicate malaria from the population as a whole (Bousema et al., 2007; Bousema et al., 2010). To date, transmission-blocking vaccines have focused on a small number of sexual stage proteins expressed on male and female gametes that play critical roles in fertilization and ookinete development in the mosquito gut (Angrisano et al., 2017). Antibodies targeting these proteins have been shown to interefere with parasite development and several (most notably, pfs25, pfs48/45, Pfs230, and HAP2) are being actively pursued as potential transmission-blocking vaccine candidates (Barr et al., 1991; Chowdhury et al., 2009; Coban et al., 2004; Farrance et al., 2011; Lee et al., 2016; Van Dijk et al., 2010; van Schaijk et al., 2006).

Despite their promise, transmission-blocking vaccines targeting sexual stage antigens face a number of challenges, one of the most critical being the large-scale, low cost manufacture of recombinant malaria proteins. While bacterial expression systems have the potential to meet that challenge, transmission-blocking vaccine candidates tend to be highly disulfide-bonded and their expression in *E. coli* and other systems is often accompanied by protein misfolding and the formation of insoluble inclusion bodies which limit their utility as vaccine antigens (Jürgen et al., 2010). To address this problem, efforts are being made to engineer *E. coli* strains capable of generating properly folded eukaryotic proteins and/or to identify solubility tags that render such proteins soluble when over-expressed in bacterial systems.

Recently, we showed that granule lattice protein 1 (Grl1p) of the free-living ciliate *Tetrahymena thermophila* substantialy improves protein solubility when expressed as a C-terminal fusion with the transmission-blocking vaccine candidate, Pfs25, in *E. coli* strains that co-express bacterial chaperone proteins (Agrawal et al., 2019). Grl1p is one of a small family of acidic proteins that populate dense core secretory granules of *T. thermophila* and that can self-assemble into nanometer-sized particles *in vivo* and *in vitro* (Cowan et al., 2005). Here, we expand on our previous study by examining a second granule lattice protein from *Tetrahymena*, Grl3p, for its ability to act as a solubility tag in the presence and absence of chaperone co-expression.

## 2. Materials and Methods

### 2.1 Materials

#### 2.1.1 *E. Coli* strains

One Shot TOP10 chemically competent *E. coli* cells (Thermofisher Scientific) were used for amplification of plasmid constructs. Protein expression studies were conducted using SHuffle® T7 Express competent *E. coli* cells (New England Biolabs, Cat No: C3029J).

#### 2.1.2 Plasmid DNA Constructs

Plasmid constructs harboring the Pfs25 or Pfs48/45 coding sequences linked to the Grl1p proprotein sequence at their 3’-ends with a TEV cleavage site between them were as previously described (Agarwal et al., 2019). For Grl3p fusion constructs, the Grl3p preproprotein sequence (accession no. <u>AF031319.1</u>) was analyzed using SignalP 5.0 (Armenteros et al., 2019) to identify the most likely cleavage site of the N-terminal signal peptide. A codon-optimized cDNA corresponding to the Grl3p proprotein (amino acids 18-377), containing TEV cleavage site at 5’ end, flanked with *Bam*HI and *Xho*I restriction sites was then prepared as a synthetic DNA fragment (Genscript Inc.). For fusion constructs containing the maltose binding protein (MBP), the coding region of MBP was modified from a pET28-MBP-TEV plasmid vector (Addgene). cDNAs corresponding to either Grl3p or the maltose binding protein were then cloned into a pET21a(+) vector with Pfs25 or Pfs48/45 in-frame and proximal to either 10X-or 6X-histidine tags for detection and purification purposes.

Plasmids for co-expression of SurA PPIase and MPD2 Foldase chaperones were obtained from New England Biolabs (Ipswich, MA, USA). All combinations of fusion constructs and chaperone proteins are listed in Supplementary Table 1.

### 2.2 Methods

#### 2.2.1 Plasmid construction

Plasmid (pUC57) DNA containing the synthetic Grl3 insert was digested with both *Bam*HI and *Xho*I enzymes. Digested DNA was run on agarose gel (1%) electrophoresis and a fragment corresponding to the Grl3p cDNA was purified by gel extraction. The fragment was then ligated to the plasmids containing either Pfs25 or Pfs48/45 as previously described (Agrawal et al., 2019).

A cDNA fragment for MBP was generated by PCR using pET28 MBP-TEV plasmid as a template and primers containing a TEV cleavage site in-frame and proximal to MBP and flanked by *Bam*HI and *Xho*I restriction sites at the 5’- and 3’-ends, respectively. Chaperone plasmids were transformed into *E. coli* TOP10 cells for amplification. Plasmids were then purified and transformed into previously prepared Shuffle T7 Express *E. coli* cells that contained either the Pfs25 or Pfs48/45 fusion constructs. Transformants were selected for double drug resistance to carbenicillin and chloramphenicol at 100µg/ml and 34µg/ml, respectively.

#### 2.2.2 Expression and purification of fusion proteins

SHuffle® T7 Express competent *E. coli* cells were transformed according to the manufacturer’s instructions. Transformants were selected for carbenicillin resistance and tested for the presence of the correct plasmid DNA by analytical digestion followed by DNA sequencing.

Expression of heterologous proteins was performed by diluting overnight cultures by a factor of 100X in 30 ml of LB media and growth at 37°C with shaking (250rpm) until they reached OD_600_=0.5. Cultures were then chilled on ice for 10 min, induced with 100 µM IPTG, and cultured at 16°C with shaking for 16 hrs. Under these conditions, cells reached an average OD of 2.0 with slight variation (±10%). For comparative analyses, cell numbers were normalized after harvesting.

For preliminary testing, cultures were centrifuged and pellets directly loaded onto duplicate SDS-PAGE gels after denaturing in SDS sample buffer containing 10% ß-mercaptoethanol at 95°C for 10min. Gels were stained with Coomassie for total protein or blotted onto nitrocellulose membranes for protein detection with anti-His antibodies as described below. For larger scale purification, cultures were harvested at the end of overnight incubation via centrifugation at 8000Xg for 10 min at 4°C. Supernatants were decanted and cell pellets frozen at -20°C for downstream processing.

##### Purification

Frozen cell pellets were thawed on ice and resuspended in 20mM Tris, 300mM NaCl, 5% glycerol, 1% Tween-80, and 15mM imidazole (pH 8.0) containing protease inhibitors (Roche cOmplete EDTA-free) as previously described (Agrawal et al., 2019). Cell suspensions were lysed by sonication (10 cycles of 10 sec on, 10 sec off at 80% power) (Qsonica Sonicator Q125, Fisher Scientific). Soluble and insoluble fractions from each cell lysate were separated by centrifugation at 20,000xg for 40min at 4°C and used to determine the percent soluble protein as described in the text.

Fusion proteins were purified using Ni-NTA resin according to manufacturer’s instructions (Thermofisher, R90115). Briefly, 1 ml of resin (2 ml 50% slurry) was poured in a 5 ml disposable column. Up to 5 ml of lysate was applied to the column followed by washing in 20 mM imidazole, and elution of recombinant fusion proteins with increasing concentrations of imidazole between 100-500 mM.

#### 2.2.3 SDS-PAGE and Western Blot analysis

Samples were mixed with Laemmli Sample Buffer (BioRad, 1610737EDU) containing 10% β-mercaptoethanol. After boiling for 5 min, samples were loaded on 12% polyacrylamide gels with a 4% stacking layer. Electrophoresis was performed in Tris-Glycine buffer at 200 V for 45 min at an average gel length of 8 cm. For Coomassie staining, gels were washed in warm water for 10 min and then stained with 0.4% Coomassie in Methanol/Acetic acid/Water (6:1:13) for 10 min followed by de-staining in 10% methanol/10% acetic acid until clear backgrounds were achieved. For western blots, gels were transferred to nitrocellulose membranes by wet transfer (2 h, 80 V, with chilled buffer). Membranes were transferred to blocking buffer containing 1.5% bovine serum albumin in PBS containing 0.1% Tween-20 and gently rocked for 1 hr at RT. Blocking buffer was refreshed and mouse anti-His antibody (Thermofisher, Cat#MA1-21315) was added at a dilution factor of 1:2000 in blocking buffer. Membranes were incubated at 4°C with gentle rocking overnight. HRP-conjugated secondary antibody (Goat Anti-Mouse Ig; Southern Biotech, Cat #:1010-05) was diluted 1:1000 in PBS containing 3% nonfat milk powder and membranes were incubated at room temperature for 1h with gentle rocking. Membranes were washed three times with PBS-Tween between each step. For signal detection, membranes were incubated in SuperSignal West Pico Chemiluminescent Substrate (Thermo Scientific) at a ratio of 1 ml per 20 cm^2^ of membrane and images captured using a BioRad gel imager.

#### 2.2.4 Analysis of protein charge structure and relative solubility

Electrical charge distributions of Grl1p and Grl3p proteins were analysed using the online protein analysis tool, ProteinSol (Hebditch et al., 2017). Following high-speed centrifugation, fractionation by SDS-PAGE and Western blotting, the relative signal strength in bands corresponding to Pfs25 or Pfs48/45 chimera in the soluble and insoluble protein fractions were quantitated using ImageJ (Rasband, 1997-2018) software.

## 3. Results

### 3.1. Culture growth, expression and purification of Grl1-tagged fusion proteins

To validate expression of Grl-fusion proteins in *E. coli*, transformed cells harboring a cDNA construct encoding malarial Pfs25 linked to Grl1p (Pfs25-TEV-Grl1-His) were cultured and induced under varying conditions of IPTG concentration, temperature and time. Cells were then lysed and resulting material separated into soluble and insoluble fractions by high-speed centrifugation. As shown in Figure 1, cells induced at 16°C produced prominent bands of ∼90 kDa that stained positively with an antibody against the C-terminal His tag in Western blots. While the majority of this protein appeared in the insoluble fraction, bands of the same size were also present in the soluble fractions. As noted previously (Agrawal et al., 2019; Chilcoat et al., 1996), Grl1p migrates anomalously on SDS-PAGE which likely accounts for the difference in the predicted molecular mass of the Pfs25-Grl1p chimera (65kDa) compared with the size determined by gel electrophoresis (∼90 kDa) (Figure 1). Varying the IPTG concentrations between 50-500 mM had only a small effect on overall expression (Figure 1), while induction at 30°C resulted in significantly lower protein expression than at 16°C. Finally, the Pfs25-Grl1p fusion could be readily purified from the soluble fraction by affinity chromatography on Ni-NTA as shown in Figure 1.

**Figure 1.**
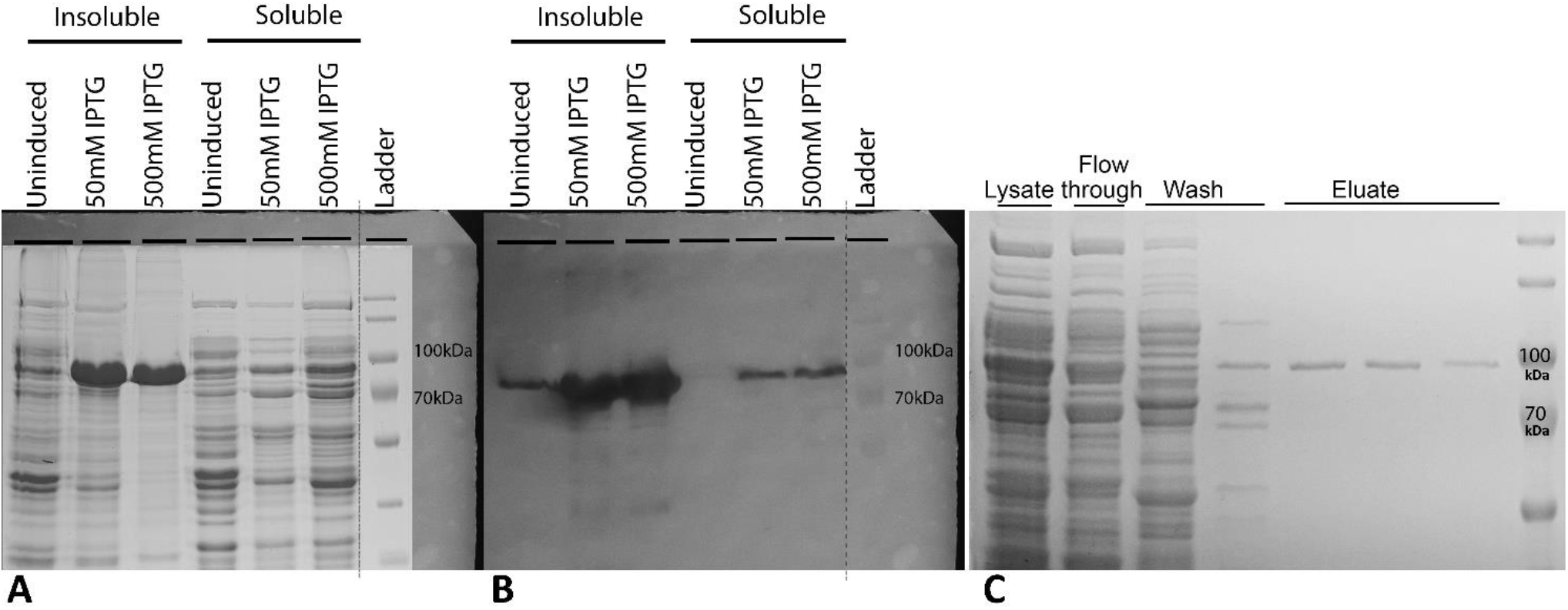
Expression of Pfs25-TEV-Grl1 fusion protein under different conditions. Shuffle T7 Express *E. coli* cells were induced with increasing concentrations of IPTG at 16°C, harvested and then lysed by sonication 4 hr after induction. Followig high-speed centrifugation, the cell pellet and soluble protein fractions were subjected to SDS-PAGE and either stained with Coomassie blue (A) or processed for Western botting using an antibody against the C-terminal His-tag (B). Note the presence of a prominent band of ∼ 90kDa in the insoluble fractions that stained heavily with anti-His antibody in the Western blot. Positively stained material was also present in the soluble fractions. When applied to Ni-NTA and eluted with increasing concentrations of imidazole (Wash: 10mM, Eluate 1: 50mM, Eluate 2: 100mM. Eluate 3: 200mM), the soluble fractions yielded a highly purified protein of the expected size of the fusion protein on SDS-PAGE (C).

### 3.2 Expression of Grl3-tagged Pfs25 and Pfs48/45 fusion proteins

In addition to Grl1p, *T. thermophila* expresses a number of other granule lattice proteins, with Grl3p being among the most abundant. As in the case of Grl1p (predicted pI: 4.28), Grl3p is also acidic (predicted pI: 4.21) and contains numerous patches enriched in aspartic and glutamic acid residues (Figure 2).

**Figure 2.**
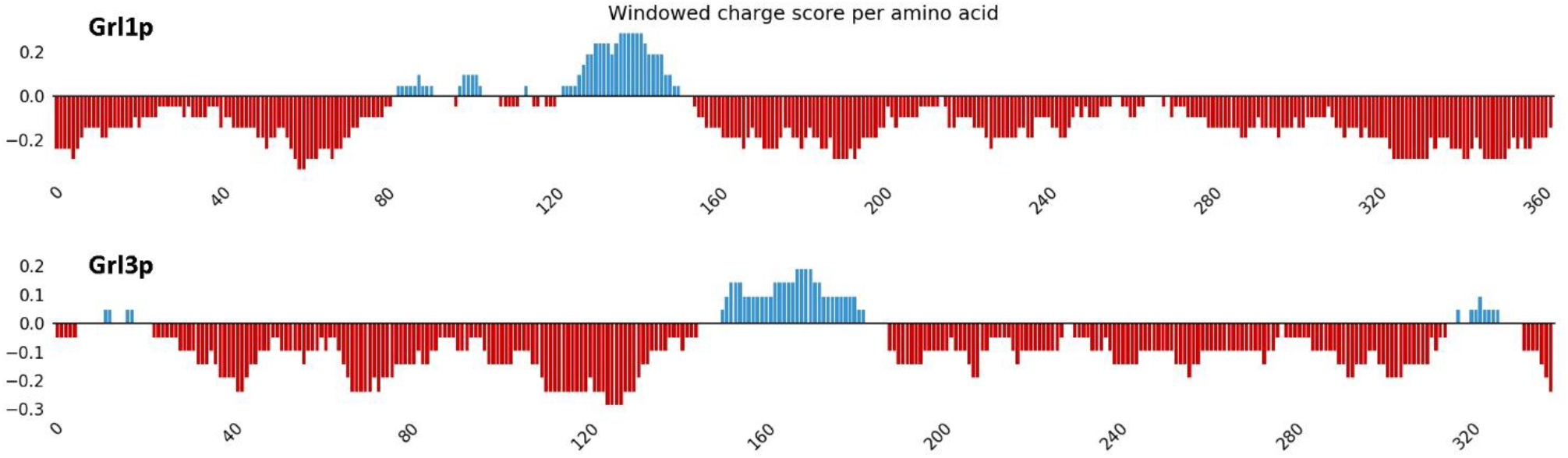
Comparison of amino acid charge distribution in Grl1p and Grl3p determined with the online tool Protein-Sol (Hebditch et al., 2017).

To determine whether malaria antigens could be expressed as chimeras with Grl3p, we placed the coding region of Grl3p downstream of either Pfs25 or Pfs48/45 with a TEV cleavage sequence in between, and introduced the resulting constructs into *E. coli* SHuffle® T7 Express.

As shown in Figure 3, transformants harboring these constructs were successfully expressed yielding both soluble and insoluble forms of the Grl3p fusion proteins. As with Grl1p chimera, highest levels of total protein production were seen at 16°C, where the metabolism of *E. coli* was slowed down to allow more time for proper protein folding (data not shown). Expression levels for the Pfs25 fusion proteins were generally higher compared to Pfs48/45 fusions although, in each case, variation between clonal replicates were noted.

**Figure. 3.**
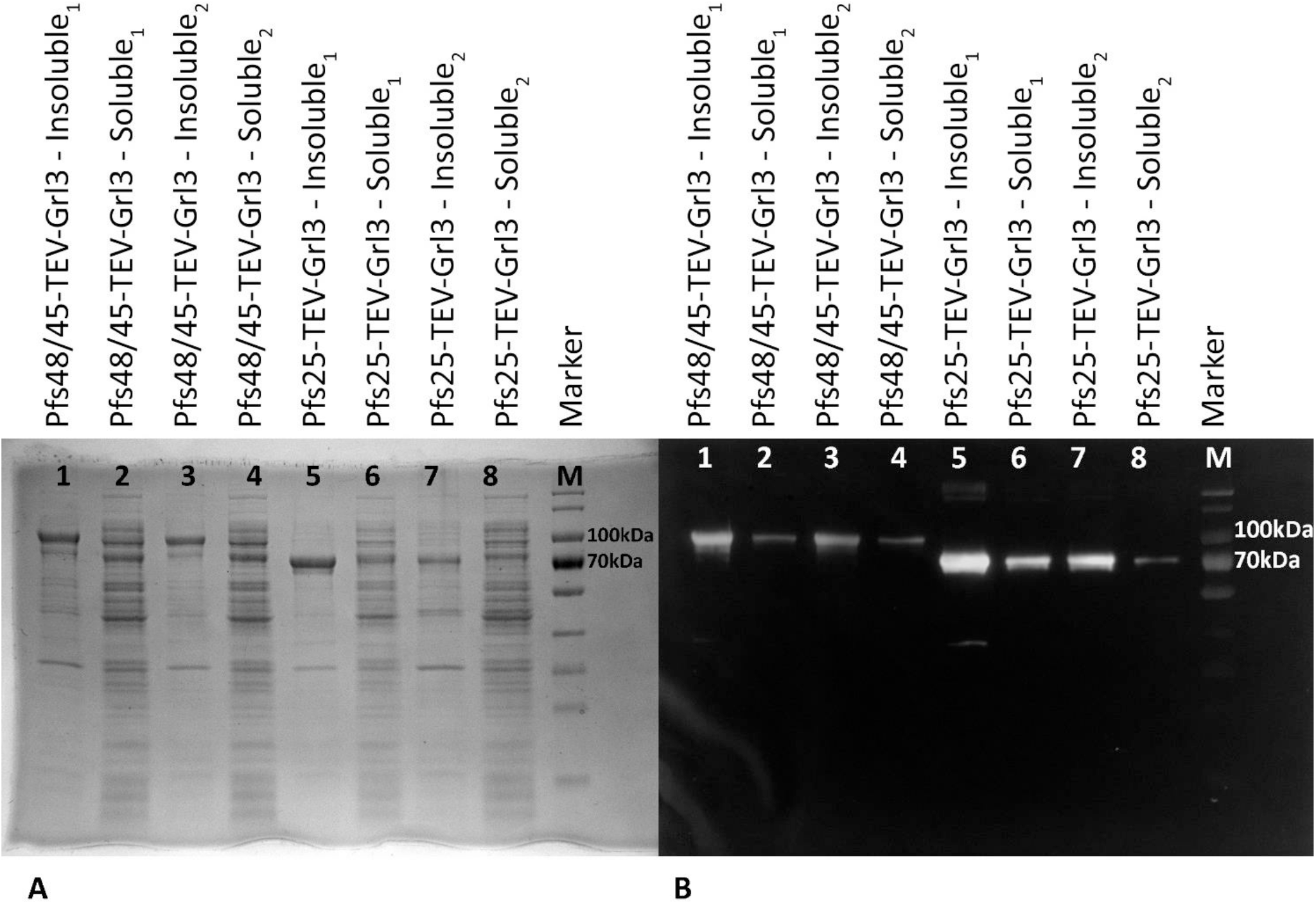
Expression of Grl3p chimeras. Left Panel: Transformants harboring his-tagged versions of either Pfs48/45-TEV-Grl3 (lanes 1-4) or Pfs25-TEV-Grl3 (lanes 5-8) constructs were cultured, induced with IPTG and subjected to high speed centrifugation. Soluble and insoluble fractions were then fractionated by SDS-PAGE and either stained with Coomassie blue (A), or processed for Western blotting using anti-his antibodies (B). Each lane is labeled with the corresponding sample loaded. Subscript numbers indicated the clonal replicates, that is, the first and second colonies chosen for expression of that protein.

While the SHuffle strain used to express these constructs was engineered to promote disulfide bond formation, we examined the effect of two different chaperone proteins, SurA PPIase and MPD2 foldase (Goemans et al., 2014; Zapun et al., 1999), on overall production of soluble and insoluble forms of Pfs25- and Pfs48/45-Grl3p chimera, respectively, following co-expression in the same cell. We then compared the results with expression of the same antigens linked to either Grl1p, or the well studied solubility tag, maltose binding protein (MBP). Table 1 shows the relative percent expression of soluble and insoluble protein for each of the chimera expressed in the presence or absence of two different chaperone proteins. Based on quantitative Western blotting, we found that the total expression of recombinant protein was similar regardless of the tag or the chaperone protein (data not shown). Nevertheless, the relative percentage of soluble and insoluble protein varied considerably depending on the antigen, the solubulity tag and the chaperone used. Most notably, when compared to MBP, the Grl1p or Grl3p tags yielded the highest percent expression of soluble protein in every case except the Pfs25-MBP fusion in combination with SurA PPase, which had a relative solubility of 19.2 %. Furthermore, the Pfs48/45 construct linked to Grl3p and co-expressed with the MDP2 foldase, showed the highest percent expression of soluble protein (23.5%) among all constructs and conditions tested (Table 1).

**Table 1.**
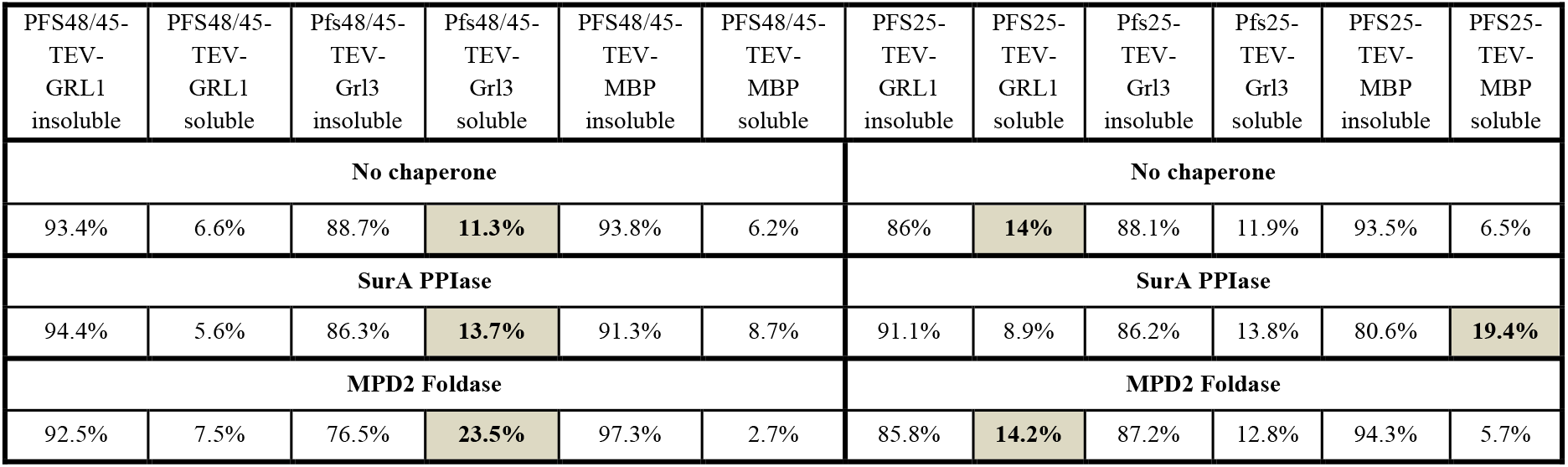
Densitometry results for all combinations of fusion proteins and chaperone plasmid. Highest percent solubility was highlighted in gray for each of 6 sets.

| PFS48/45-TEV-GRL1<br>insoluble | PFS48/45-TEV-GRL1<br>soluble | Pfs48/45-TEV-GrI3<br>insoluble | Pfs48/45-TEV-GrI3<br>soluble | PFS48/45-TEV-MBP<br>insoluble | PFS48/45-TEV-MBP<br>soluble | PFS25-TEV-GRL1<br>insoluble | PFS25-TEV-GRL1<br>soluble | Pfs25-TEV-GrI3<br>insoluble | Pfs25-TEV-GrI3<br>soluble | PFS25-TEV-MBP<br>insoluble | PFS25-TEV-MBP<br>soluble |
| --- | --- | --- | --- | --- | --- | --- | --- | --- | --- | --- | --- |
| No chaperone |  |  |  |  |  | No chaperone |  |  |  |  |  |
| 93.4% | 6.6% | 88.7% | 11.3% | 93.8% | 6.2% | 86% | 14% | 88.1% | 11.9% | 93.5% | 6.5% |
| SurA PPIase |  |  |  |  |  | SurA PPIase |  |  |  |  |  |
| 94.4% | 5.6% | 86.3% | 13.7% | 91.3% | 8.7% | 91.1% | 8.9% | 86.2% | 13.8% | 80.6% | 19.4% |
| MPD2 Foldase |  |  |  |  |  | MPD2 Foldase |  |  |  |  |  |
| 92.5% | 7.5% | 76.5% | 23.5% | 97.3% | 2.7% | 85.8% | 14.2% | 87.2% | 12.8% | 94.3% | 5.7% |

## 4. Discussion

One of the key hurdles for the production of malaria vaccine antigens in *E. coli* has been the low solubility of recombinantly expressed proteins. This extends to important transmission-blocking vaccine candidates such as Pfs25 and Pfs48/45 (Nikolaeva et al., 2015), and while protein denaturation and refolding offers a potential solution to the problem (Singh and Panda, 2005), renaturation of highly disulfide bonded malaria proteins is inefficient and adds significant cost to the manufacturing process.

The use of *E. coli* strains that have been modified to promote disulfide-bond formation and protein folding in the bacterial cytosol offer an alternative approach towards improving the solubility of heterologously expressed proteins (Berkmen et al., 2005; Gurmu et al., 2009; Lobstein et al., 2012; Locker and Griffiths, 1999). *E. coli* SHuffle represents one such strain and was specifically designed to increase disulfide bonding through constitutive expression of a modified disulfide bond isomerase (Dsbc) in the *trxB gor* suppressor strain, SMG96 (Lobstein et al., 2012).

Under appropriate conditions, *E. coli* SHuffle has shown considerable success in increasing the solubility of a variety of eukaryotic proteins compared with expression in standard *E. coi* K12 strains (Ke and Berkmen, 2014; Lobstein et al., 2012; Nozach et al., 2013). Indeed, by combining expression in SHuffle with co-expression of selected chaperones and/or the use of solubility tags, further improvements in solubility can be achieved (Bessette et al., 1999; Jurado et al., 2002; Yuan et al., 2004). Notably, this combined approach was effective in generating soluble forms of the malaria transmission-blocking vaccine, Pfs25, as a chimera with the granule-lattice protein, Grl1p, from *Tetrahymena thermophila* (Agrawal et al., 2019). In the absence of tags, soluble Pfs25 was barely detectable even in the SHuffle strain, while Pfs48/45 appeared to be completely insoluble (Agrawal et al., 2019). This is entirely consistent with previous studies (Gregory et al., 2012; Kumar et al., 2014; Nikolaeva et al., 2015) and the results obtained here.

The granule lattice proteins of *T. thermophila* assemble into crystalline arrays of densely packed nanoparticles that could serve as vehicles for the delivery of recombinant subunit vaccines (Jayaram et al., 2010). While it is unclear whether Grl3p can form higher order structures either alone or in combination with vaccine antigens, we wanted to test its relative ability to act as a solubility tag in comparison with Grl1p and the well-studied, commercially available MBP tag. Like Grl1p, Grl3p is highly acidic and has a similar overall charge distribution (Figure 2) suggesting that the biophysical properties of the two proteins are similar. Indeed, Grl3p clearly mimicked the ability of Grl1p to generate soluble forms of Pfs25, and importantly, generated the highest levels of soluble Pfs48/45 for any of the tags under the conditions tested (Table 1). Finally, different combinations of tags and chaperones produced different yields of soluble protein and while the MBP tag in combination with SurAPPase had the highest relative solubility for Pfs25, in the case of Pfs48/45, the Grl3 tag in combination with MPD2 Foldase produced almost a nine-fold increase in solubility compared to the MBP tag paired with the same chaperone. Such differences argue for the possibility of synergistic effects between chaperones and tags which necessitate the testing of all possible combinations to determine optimal yields of soluble protein.

In conclusion, we demonstrate that Grl3p and Grl1p are viable solubility tags for proteins that are difficult to express in soluble form in *E. coli*. As expected, the use of additional chaperone expressing plasmids further improved solubility in the *E. coli* Shuffle T7 Express strain. Continued advances along these lines are expected to enhance expression of *Plasmodium* antigens in *E. coli* and further the development of transmission-blocking vaccines for malaria.

## Acknowledgments

We thank Dr. Alka Agrawal for providing backbone plasmids for Grl1 and Pfs fusion proteins as well as suggestions on soluble protein analysis. We also thank Dr. Mehmet Berkmen for his guidance in the use of chaperone plasmids. We thank Mr. Daniel Kolbin for his helpful discussions of the data and Mr. Mozammal Hossein for his excellent technical support on the project.

## Author Contributions

C.A. performed the major experiments and wrote the manuscript. D.M.C.H. helped in the design of experiments and data interpretation. T.G.C. was involved in designing the research, data interpretation and writing/editing of the manuscript.

