## Supplementary Table 1 for "*Tetrahymena thermophila* Granule Lattice Protein 3 Improves Solubility of Sexual Stage Malaria Antigens Expressed in *Escherichia coli*"

**Supplementary material**

**Supplementary Table 1.** List of the constructs used to express Pfs25, Pfs48/45 and their fusions with either Grl1p, Grl3p or MBP. All fusion proteins were also co-expressed either with SurA Pplase or MPD2 Foldase in separate experiments.

| Pfs25 constructs | Pfs48/45 constructs |
| --- | --- |
| pET21a(+)Pfs25 | pET21a(+)Pfs48/45 |
| pET21a(+)Pfs25-TEV-Grl1 | pET21a(+)Pfs48/45-TEV-Grl1 |
| pET21a(+)Pfs25-TEV-Grl3 | pET21a(+)Pfs48/45-TEV-Grl13 |
| pET21a(+)Pfs25-TEV-MBP | pET21a(+)Pfs48/45-TEV-MBP |
